# IDO-1 inhibition improves outcome after fluid percussion injury in adult male rats

**DOI:** 10.1101/2023.10.31.564953

**Authors:** Marawan Sadek, Kurt R Stover, Xiaojing Liu, Mark A Reed, Donald F Weaver, Aylin Y Reid

## Abstract

The enzyme indoleamine 2,3 dioxygenase 1 (IDO1) catalyzes the rate-limiting step in the kynurenine pathway (KP) which produces both neuroprotective and neurotoxic metabolites. Neuroinflammatory signals produced as a result of pathological conditions can increase production of IDO1 and boost its enzymatic capacity. IDO1 and the KP have been implicated in behavioral recovery after human traumatic brain injury (TBI), but their roles in experimental models of TBI are for the most part unknown. We hypothesized there is an increase in KP activity in the fluid percussion injury (FPI) model of TBI, and that administration of an IDO1 inhibitor will improve neurological recovery. In this study adult male Sprague-Dawley rats were subjected to FPI or sham injury and received twice-daily oral administration of the IDO1 inhibitor PF-06840003 (100 mg/kg) or vehicle control. FPI resulted in a significant increase in KP activity, as demonstrated by an increased ratio of kynurenine:tryptophan, in the perilesional neocortex and ipsilateral hippocampus three days post-injury (DPI), which normalized by seven DPI. The increase in KP activity was prevented by PF-06840003. IDO1 inhibition also improved memory performance as assessed in the Barnes maze and anxiety behaviors as assessed in open field testing in the first 28 DPI. These results suggest increased KP activity after FPI may mediate neurological dysfunction, and IDO1 inhibition should be further investigated as a potential therapeutic target to improve recovery.

**Significance Statement:** The kynurenine pathway and its rate-limiting enzyme indoleamine 2,3 dioxygenase 1 (IDO1) have been implicated in a variety of neurological disorders, including traumatic brain injury. We have demonstrated increased IDO1 activity in male rats after fluid percussion injury, a widely used model of traumatic brain injury. Pharmacological IDO1 inhibition after fluid percussion injury improved performance on tests of memory and anxiety-like behaviors, demonstrating a role for IDO1 in traumatic brain injury outcomes and supporting further investigation into its potential as a therapeutic target.

## Introduction

Under normal conditions, the kynurenine pathway (KP) is the primary route for tryptophan catabolism in neuronal support cells (Savitz, 2020). This pathway is essential in the proper synthesis of tryptophan and is important in the promotion of neuroregulatory pathways (Badawy, 2020; Savitz, 2020). The KP is regulated by the enzymes tryptophan 2,3-dioxygenase (TDO2) and indoleamine 2,3 dioxygenase 1 (IDO1) (Hattori et al, 2018; Pallotta et al, 2021). IDO1 appears to be the key enzyme regulating the KP in pathological conditions (Yan et al, 2015; Pallotta et al, 2021). In the brain, IDO1 is expressed in microglial, neuronal, and astroglial cells and metabolizes free-form tryptophan that has crossed the blood-brain barrier into several metabolites, most notably neuroprotective kynurenic acid and neurotoxic quinolinic acid (QuinA) (Yan et al, 2015). Overactivation of IDO1 has been directly linked with increases in neuroinflammation in several diseases and to long-term memory dysfunctionality, motor deficits and mood disorders such as depression and anxiety (Dobos et al., 2012; Mazarei & Leavitt, 2015; Wu et al., 2013). There has been a recent interest in the role of the KP in consequences of traumatic brain injury (TBI), with higher levels of neurotoxic QuinA in the CSF and/or plasma of humans with TBI correlating with poorer recovery (Yan et al., 2015; Singh et al., 2016; Meier & Savitz, 2022) and increased mortality (Sinz et al., 1998). There is sparse information about the KP in experimental TBI. Rodent (Chung et al., 2009) and rabbit (Zhang et al, 2018) models of TBI have demonstrated increased IDO1 activity or QuinA levels, but IDO1 activity has not been investigated in commonly used TBI models such as fluid percussion injury (FPI). Furthermore, it is unknown whether manipulation of the KP after TBI impacts functional outcome.

IDO1 inhibition is being investigated for its beneficial effects in a number of disorders, with the majority of research to date focusing on cancer. Most studies of pharmacological IDO1 inhibition in cancer have reported an overall improvement in health with decreased tumor formation, less cellular metastases, or increased subject survival rate (Ahlstedt et al., 2020; Lynch et al., 2020; Tian et al., 2019). Of the commercially available IDO1 inhibitors, the drug PF-06840003 is a highly selective IDO1 antagonist with a half-life of 16-19 hours in rats and mice (Crosignani et al., 2017), reducing kynurenine levels and IDO1 activity (Reardon et al., 2020; Gomes et al., 2018; Liu et al., 2021). PF-06840003 has already been shown to have promising effects in a clinical trial for glioma treatment (Reardon et al., 2020). Overall, multiple lines of evidence demonstrate PF-06840003 safely diminishes IDO1 activity in cancer, which may also prove beneficial in the treatment of neurological disorders. If IDO1 inhibition can limit IDO1 activity in response to TBI, it may result in improved outcomes.

In this study, we orally administered PF-06840003 to adult male Sprague-Dawley rats after FPI or sham injury. We determined the effect of TBI and IDO1 inhibition on KP activity and assessed functional outcomes in a variety of neurological domains.

## Materials and Methods

### Experimental Subjects and Study Design

This study used young adult male Sprague-Dawley rats (Charles River, Canada) weighing 229-328g. All rats were outbred at the animal breeding facility and arrived at the local animal care facility one week before the procedures. Animals had free access to food and water and were handled daily for one week prior to any procedures. Rats were randomly assigned to one of four groups: Sham+Vehicle, Sham+Inhibitor, TBI+Vehicle, or TBI+Inhibitor. Group assignments were made by a research assistant who was not involved in any data collection or analyses and kept coded until the completion of all analyses. Experimenters were blind to the identity of the groups during the entirety of the experiment. As both sham and TBI groups underwent craniotomy and subsequent suturing of the scalp, these groups were not visually distinguishable. Rats underwent either FPI or sham injury and then received an IDO1 inhibitor, PF06840003 (100 mg/kg), or an equivalent volume of vehicle ((2-Hydroxypropyl)-β-cyclodextrin (HBCD)), by oral gavage twice daily. For blinding of drug treatment, syringes containing experimental drugs or vehicle were prepared by the research assistant, labelled with a coded ID, and covered with an opaque material so the contents were not visible to the investigators administering the compounds. One subset of rats was sacrificed at three- or seven-days post-injury (DPI) and the ratio of kynurenine:tryptophan (KT) was measured as a marker of KP activity. A second subset of animals received vehicle or treatment for 28 days, during which they were evaluated on their motor ability and coordination on the composite Neuroscore test; their visuospatial learning and memory on the Barnes maze test; as well as their anxiety behaviors in the open field test. KT ratios were measured in these rats after behavioral testing was complete.

### Ethical Approval

Experiments were performed in accordance with the policies and guidelines established by the Canadian Council on Animal Care and ethical approval from the University Health Network-Animal Resources Centre.

### Drug Pharmacokinetics

There are no reports of the use of systemic PF-06840003 administration for central nervous system disorders in rats. Cancer studies using this inhibitor have typically used mouse models with oral administration of the drug at doses that range from 20 to 60mg/kg daily with some studies administering the drug interperitoneally in mice at a dose of 200mg/kg (Dolšak et al., 2021; Cheong et al., 2018; Gomes et al., 2018, Liu et al., 2021). In order to determine appropriate dosing for this study, initial pharmacokinetic studies were performed in rats with PF-06840003 (10 mg/kg, oral gavage) demonstrating good brain exposure and a half-life of ∼3.5 hours. Sprague-Dawley rats (n=3/time point) were dosed with compound via oral gavage at 10mg/kg and then sacrificed at one of the seven scheduled time points (0.25, 0.5, 1, 2 5, 8 or 24 h) post dose. The concentration of brain and plasma PF-06840003 was determined via LC/MSMS. With half maximal effective concentration EC50 (hIDO)=0.5 μM (∼120 ng/mL), and evidence that maximal IDO1 inhibition is achieved at greater than four times the EC50, the goal free fraction was >500 ng/mL. Based on estimates of the percent of unbound drug from previous mouse data, we calculated the dose needed to achieve 500 ng/mL at 12 hours (timing of the subsequent dose) was at least 50 mg/kg. Based on these data, we proceeded to a drug tolerability study using 100 mg/kg twice daily. PF-06840003 was synthesized according to previously published protocols (Crosignani et al, 2017).

### Drug Tolerability Study

Naïve adult Sprague-Dawley rats were used for drug tolerability studies. Vehicle solution (HBCD, 20% w/v dissolved in water; Sigma-Aldrich #332593) or IDO1 inhibitor (10 mg of PF-06840003/ml of HBCD solution) were administered at a dose of 100mg/kg by oral gavage every twelve hours for 14 days. Rat weight, grooming, movement and feeding behaviors were monitored throughout the study. Rats were also examined for any signs of discharge, distress, or discomfort.

### Fluid Percussion Injury

Rats were anesthetized with 4% isoflurane, the head was shaven, and the scalp cleaned with an antiseptic solution. A midline incision was made through the scalp to expose the skull and a craniotomy was performed by drilling a hole with a radius of 2.5 mm, positioned 1mm posterior to bregma and 3mm lateral to the midline over the left parietal neocortex (Reid et al., 2016; Casillas-Espinosa et al., 2019; Ndode-Ekane et al., 2019; Saletti et al, 2023). Care was made not to penetrate the dura while drilling or removing the bone flap. A plastic injury cap was affixed over the opening with dental cement and then filled with saline. The rats were then removed from isoflurane and monitored for return of the toe-pinch reflex. As soon as this reflex was present, rats were moved to a platform where the injury cap was directly connected to the central injury shaft of the FPI device (Virginia Commonwealth University). A standard FPI device was used with a 63 cm long glass cylinder, with a diameter of six centimeters, that was affixed with a moving piston on one end and an output to a tube called the central injury shaft at the other end. The pendular hammer was set to fall at an angle of 12.5 degrees, resulting in a pressure pulse duration of 10ms and a force of 3.2-3.5 atm as measured using a pressure transducer connected to an oscilloscope. The duration of apnea was measured and recorded from the moment of impact. Following the return of respiration, the animals were placed on a heating pad and timed for the return of the righting reflex. After the return of the righting reflex, the rats were placed back in a stereotaxic frame with light anesthesia for removal of the plastic injury cap and suturing of the skin of the scalp. After surgery rats were placed in a clean cage on a heating pad until they regained full mobility and were eating and drinking. They were then returned to the vivarium and monitored daily for signs of abnormal grooming, feeding or weight loss.

### Drug Preparation and Administration for Behavioral Studies

Vehicle solution consisted of (2-Hydroxypropyl)-βcyclodextrin (HBCD) powder dissolved in MilliQ water (0.2g/mL). The IDO1 inhibitor was prepared by dissolving PF-06840003 in the HBCD solution (10g/mL). All batches of PF-06840003 and HBCD were produced a maximum of three days before administration. For every 100 g of rat body weight, 1 mL of HBCD or PF-06840003 solution (100 mg/kg) was given through an oral gavage syringe attached to a 75mm flexible plastic feeding tube. Drugs were administered every twelve hours starting immediately after recovery from FPI and for the duration of the study (28 days).

### Composite Neuroscore

The composite Neuroscore was used to assess motor ability and reflexes. This test has previously demonstrated motor deficits for up to one-month post-TBI (McIntosh et al., 1989; Maegele et al., 2005; Zhang et al., 2005; Febinger et al., 2016). There were seven main parameters that contributed to the composite Neuroscore: right and left forelimb flexion, right and left hindlimb flexion, right and left resistance to lateral pulsion, and performance on the angle board. The maximum score that a rat could receive on the Neuroscore was 28, with each of the parameters scored out of four. The test was conducted at three days pre-injury to determine baseline scoring and then again at 2DPI, 8DPI, 15DPI, and 28DPI.

### Barnes Maze

The Barnes Maze was used to assess rats’ learning ability and memory of visuospatial information after TBI (Harrison et al., 2006; Kovesdi et al., 2012; Maegele et al., 2005; Rowe et al., 2014). The maze consists of a large spherical table with a diameter of 120 cm with 20 holes surrounding the edge of the table. Each hole can be blocked with a sliding blockade and one hole can hold an escape chamber. Five white walls surrounded the Barnes maze table, four of these walls were marked with a unique symbol with a unique color, while the fifth wall (entrance to the area) was left blank. Four of the walls were also affixed with bright lights that served as aversive stimuli for the rats. To test learning ability, performance on the maze was assessed on 14-17 DPI. On the first trial on 14DPI each rat was directly placed in the escape chamber connected to the target escape hole and kept there for one minute to acclimate the rats to the escape chamber outside of testing conditions. Rats were then placed in the center of the maze and allowed to roam freely for five minutes. Once a rat had located and entered the escape chamber, they were left there for one minute before the conclusion of the trial. If the rat was unsuccessful at locating the escape chamber by the end of the five-minute interval, they were guided towards it by the experimenter and then left there for one minute. Each rat completed two five-minute trials on each of the learning days and all the trials were separated by five-minute cleaning sessions using 70% alcohol.

To test short-term and long-term memory, performance on the maze was assessed in probe trials on 18DPI and 28DPI. On these days the escape chamber was removed, and the target hole was fitted with a blockade identical to the other holes around the maze. Rats were allowed to roam freely for a complete five-minute interval. Success or failure at finding the target hole did not impact the duration of the task and rats were not guided to the target hole following the five-minute interval. During these memory testing days, each rat completed only one trial each day.

### Open Field Test

The open field test (OFT) was used to assess anxiety behavior following TBI (Rowe et al., 2014). The test was conducted in a white plastic square box with the dimensions 120×120×40 cm. The box was placed in an area surrounded by five white walls, with four of those walls affixed with bright white lights. A video camera was positioned at the top and center of the area, capturing the entirety of the OFT box. Rats were monitored for two five-minute intervals for presence in the inner or outer regions of the OFT box. The inner region of the OFT box was defined as the center 36% of the area of the box with the dimensions 72×72 cm, while the outer region was defined as the parameter around the inner region.

### Brain Collection

Rats were sacrificed after the completion of the 28-day behavioral test period for perilesional cortex and hippocampal collection. Rats were anesthetized using isoflurane gas in an anesthetic chamber and then underwent trans-cardiac perfusion with 200 mL of phosphate buffer saline. Ipsilateral perilesional neocortical and hippocampal samples were extracted, flash-frozen in liquid nitrogen, and stored at -80°C.

### LCMS Analysis of the KT-Ratio

Brain samples were defrosted on ice, 1.5mL of water was added per gram of tissue, and then samples were homogenized using a handheld 1.5mL microtubule pestle. 40 µL from each homogenate was further diluted with an additional 40 µL of water, and this sample was used for LC-MS/MS analysis. The concentrations of kynurenine and tryptophan were determined via LC-MS/MS using kynurenine-13C4,15N and tryptophan-d5 as internal standards. Brain kynurenine, tryptophan and their internal standards were extracted by protein precipitation. The homogenized brain sample was protein precipitated with four volumes of 0.1 N trichloroacetic acid. Following protein precipitation, the sample mixture was centrifuged at 4000 rpm for 10 minutes at 4 °C and the supernatant was collected for analysis. These data were used to calculate the ratio of kynurenine to tryptophan present in each sample as a marker of IDO1 activity.

### Statistical Analyses

Power calculations were performed in advance to determine the sample size needed to demonstrate an effect in the behavioral tests with an α-error probability of 0.05 and a power (1-β error probability) of 0.95 using previous FPI data from our laboratory and the literature. This calculation was done using G*Power version 3.1.9.7 software for windows. *A priori* sample size calculations were run using Wilcoxon-Mann-Whitney tests. For both the composite Neuroscore and Barnes maze, at least seven rats were needed to establish an effect size of 1.940. We had not previously worked with the OFT and used data from another FPI study (Learoyd & Lifshitz, 2012) to calculate a sample size of at least eight rats per group. We therefore aimed for a minimum of 12 rats/group.

Data were analyzed using the Graphpad Prism Software version 9.4.1 for Windows (GraphPad Software, La Jolla California USA, www.graphpad.com). Differences in apnea duration and return of the righting reflex between the two TBI groups were compared using two-tailed unpaired t-tests. Repeated measures two-way analysis of variance (ANOVA) with Fisher’s LSD multiple comparisons test were used in the analyses of data from the composite Neuroscore and Barnes maze learning parameters. An ordinary one-way ANOVA with Fisher’s LSD multiple comparisons test was used in the analysis of data from the 18-day and 28-day memory probe trials of the Barnes maze test, for the 1 DPI and 25 DPI OFT data, and for kynurenine:tryptophan ratios at 3, 7, and 28 DPI. Statistical significance was set at *p*<0.05.

## Results

### FPI and sham surgery

143 animals were used in these experiments:

1. Six naïve rats were used for the drug tolerability study (three with PF-06840003 and three with HBCD).
2. 38 rats underwent sham surgery. Four sham rats were excluded for having an exceptionally poor performance on two or more behavioural tests (2DPI Neuroscore <20 and poor performance on the Barnes Maze), indicating injury to the underlying cortex. Nine sham rats were used to determine KT ratios at three DPI (five Sham+Vehicle, four Sham+Inhibitor) and 11 to determine KT ratios at seven DPI (six Sham+Vehicle, five Sham+Inhibitor). The remaining 14 rats were used in the behavioural studies (seven Sham+Vehicle, seven Sham+Inhibitor). There were no significant differences between the Sham+Vehicle and Sham+Inhibitor groups for any of the outcomes measured, therefore the two were combined into a single Sham group for all analyses.
3. 75 rats underwent FPI. 19 died (25%) due to prolonged apnea after impact, in keeping with mortality rates in this model (Kabadi, 2010). Of the remaining 56 FPI rats, 10 were excluded due to the resulting injury being too mild based on a righting reflex of ≤ 10 mins or a drop in the 2DPI Neuroscore of less than five points from baseline. Of the 46 FPI rats included in this study, 11 were used to determine KT ratios at three DPI (five TBI+Vehicle, six TBI+Inhibitor) and 10 were used to determine seven DPI KT ratios (five TBI+Vehicle, five TBI+Inhibitor). The remaining 25 rats were used for behavioural studies (12 TBI+Vehicle, 13 TBI+Inhibitor).

### Drug Tolerability

Three rats received oral HBCD solution as a vehicle treatment and three received oral PF-06840003 (100 mg/kg) twice daily for 14 days. There were no differences in the rate of weight gain between the two groups over this time period and all rats maintained proper grooming, motility, and feeding behaviours. Therefore, we proceeded with this dose of 100 mg/kg PF-6840003 for the remainder of the studies.

### Injury Parameters

The angle of the pendulum drop from the FPI device was set to 13 degrees, resulting in a pressure pulse duration of 10ms and a force of 3.2-3.5 atm. The average duration of apnea (± SD) for the TBI+Vehicle group was 49.6 +/- 58.2 sec and the average time for the return of the righting reflex was 1231.0 ± 418.9 sec. The average duration of apnea for the TBI+Inhibitor group was 35.1 ± 53.3 sec with an average return of the righting reflex of 999.8 ± 291.7 sec. There were no statistical differences between the two TBI groups for apnea duration (*t*=0.06259, *df*=21, *p*=0.54) or return of righting reflex (*t*=1.546, *df*=21, *p*=0.14).

### Kynurenine Pathway Activity After TBI

The KT ratio was measured as an indicator of KP activity in the ipsilateral perilesional neocortex and hippocampus at three and seven DPI. There was a significant difference between groups for the KT ratio in the neocortex at three DPI (*F_2,17_*=17.47, *p*<0.0001), with a significantly higher KT ratio in the TBI+Vehicle group compared with the Sham (*p*=0.0003) and TBI+Inhibitor (*p*=0.0001) groups (Fig. 1A). There was also a significant difference between groups for the KT ratio in the hippocampus at three DPI (*F_2,16_*=4.544*, p*=0.0274), with a significantly higher KT ratio in the TBI+Vehicle versus the Sham (*p*=0.0147) and TBI+Inhibitor (*p*=0.0196) groups (Fig. 1B). There were no significant differences between any groups in the neocortex (*F_2,17_*=0.1691, *p*=0.8458) or hippocampus (*F_2,17_*=0.9230, *p*=0.4163) at seven DPI (Fig. 1C, D).

**Figure 1.**
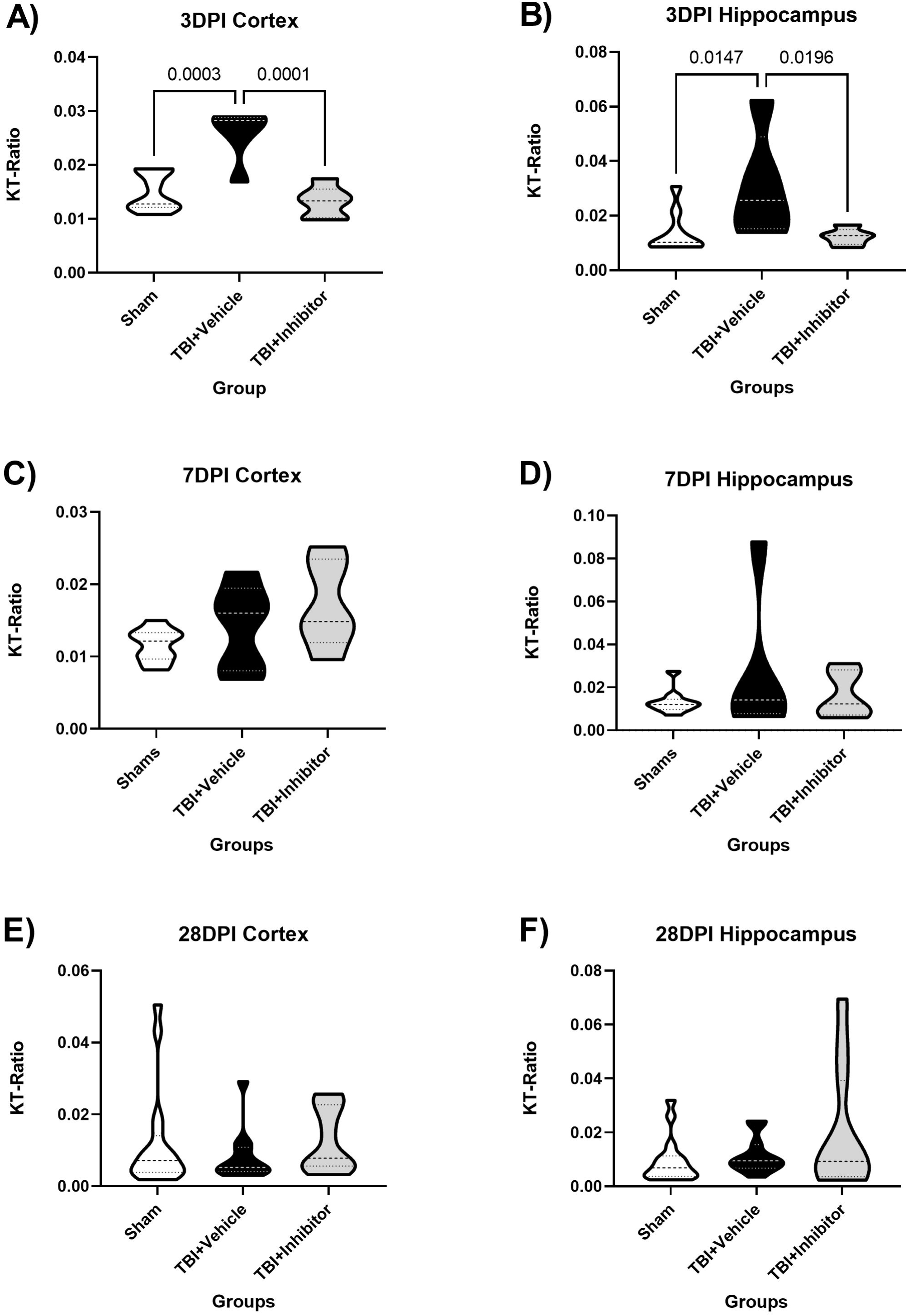
Violin plots of kynurenine:tryptophan (KT) ratios from perilesional and hippocampal brain samples after sham or fluid percussion injury. The TBI+Vehicle group had significantly higher KT ratios in samples taken from the perilesional neocortex (A) and ipsilateral hippocampus (B) three days post-injury (DPI). No differences were found between groups in either brain region at seven DPI (C, D) or 28DPI (E, F). Statistical comparisons were made with an ordinary one-way ANOVA and Fisher’s LSD at each time point and for each region.

### Neuroscore

All rats achieved a full Neuroscore of 28 during baseline pre-injury testing. Two-way ANOVA revealed a significant effect of time (*F_2.3,83.2_* =99.3; *p*<0.0001), condition group (*F_2,36_* =3.9; p<0.05) and a significant interaction (*F_8,144_* =3.51, *p*=0.001). The TBI+Vehicle group had significantly lower Neuroscores than the Sham group at 2DPI (*p*=0.017) and 8DPI (*p*=0.040) and continued to perform significantly worse compared to baseline on all testing days through to and including D28 (*p*=0.005). The TBI+Inhibitor group on the other hand only had a significantly worse performance than the Sham group at 2DPI (*p*=0.0107) and was no longer different from its baseline at 28 DPI (*p*=0.0667) (Fig. 2).

**Figure 2.**
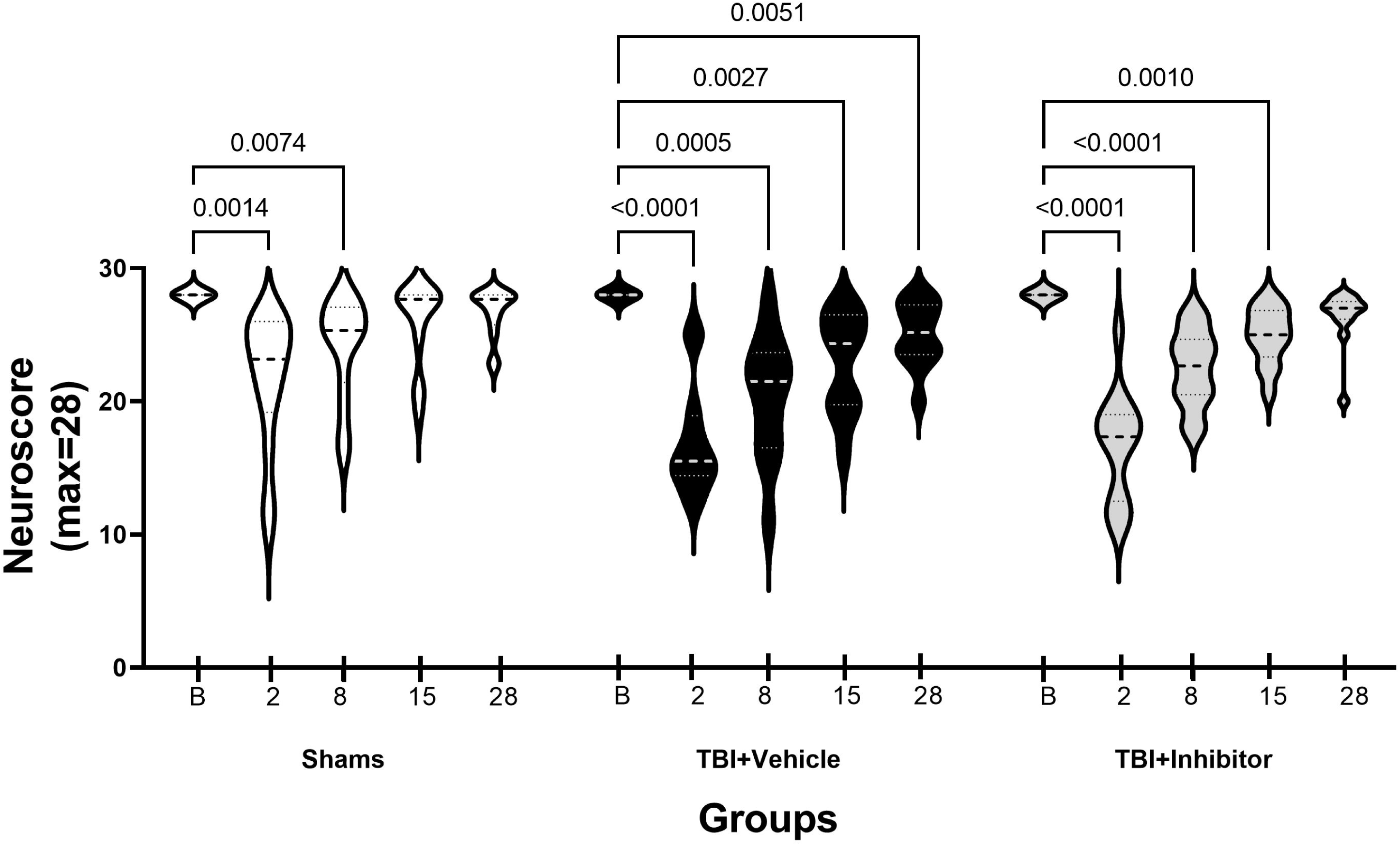
Violin plot of rat performance on the composite Neuroscore from baseline to 28 days post-injury (DPI). After fluid percussion injury, the TBI+Vehicle group performed worse than baseline on all testing days. The TBI+Inhibitor groups performed worse than baseline after injury through to 15DPI but was no longer different from baseline at 28DPI. Statistical comparisons were made using a two-wave repeated measure ANOVA and Fisher’s LSD.

### Barnes Maze Learning Phase

All groups fully learned the Barnes Maze over the four-day learning period between 14 and 17DPI (Fig. 3). Repeated measures two-way ANOVA revealed a significant effect of time on latency to locate the target hole (*F_1.9,61.96_* =24.21; *p*<0.0001) and distance travelled to the target hole (*F_1.98,61.48_*= 26.14, *p*<0.0001), with all three groups performing best on the last day of the learning phase (*p*<0.002). There were no significant differences in average latency, distance travelled, or number of errors between any of the three groups during the learning phase.

**Figure 3.**
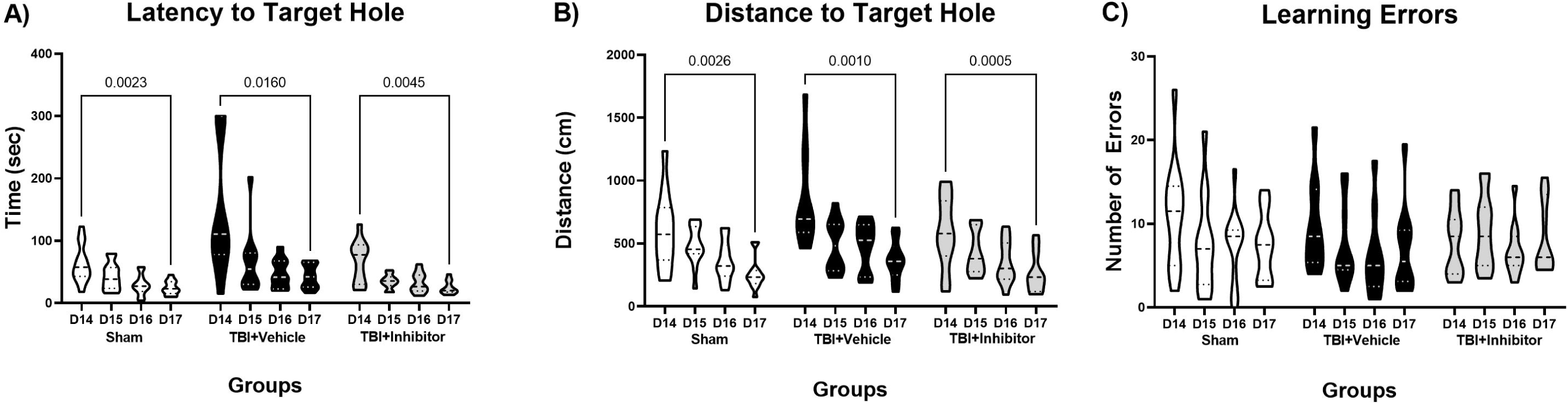
Violin plots of performance on the learning phase of the Barnes maze on days 14 through 17 post-injury. All groups full learned the Barnes maze with significant improvement over time in the latency to locate the target hole (A) and distance traveled to the target hole (B). There were no significant differences in the number of errors made over time (C), nor any differences between groups for any of the parameters. Statistical comparisons were made with two-way repeated measure ANOVAs.

### Barnes Maze Short and Long-Term Memory Probes

A short-term memory probe trial was conducted on 18DPI (Fig 4). One-way ANOVA showed a significant difference between groups in the number of errors (*F_2,30_*=3.898, *p*=0.03). Post-hoc analysis revealed the TBI+Vehicle group made significantly more errors than Sham rats before finding the target hole (*p*<0.01), while no such difference was seen between the TBI+Inhibitor and Sham group. Although not statistically significant, there was also a trend of the TBI+Vehicle group having worse performance in the other parameters, with a longer latency to find the target (*p*=0.0658). The long-term memory probe trial occurred on 28DPI (Fig 4). One-way ANOVA revealed a significant difference between groups for the latency to locate the target hole (*F_2,31_* =3.48; *p*=0.04), primary distance travelled prior to locating the target hole (*F_2,31_*=4.02; *p*<0.03), and time in the target quadrant (*F_2,31_*=4.94; *p*=0.01). Post-hoc analyses showed the TBI+Vehicle had a significantly longer latency to the target hole (*p*<0.03), travelled a significantly longer distance to locate the target hole (*p*<0.03), and spent significantly less time in the target quadrant than both the sham and TBI+Inhibitor groups (*p*<0.03). The TBI+Vehicle also performed significantly more errors compared with the TBI+Inhibitor group (*p=*0.043).

**Figure 4.**
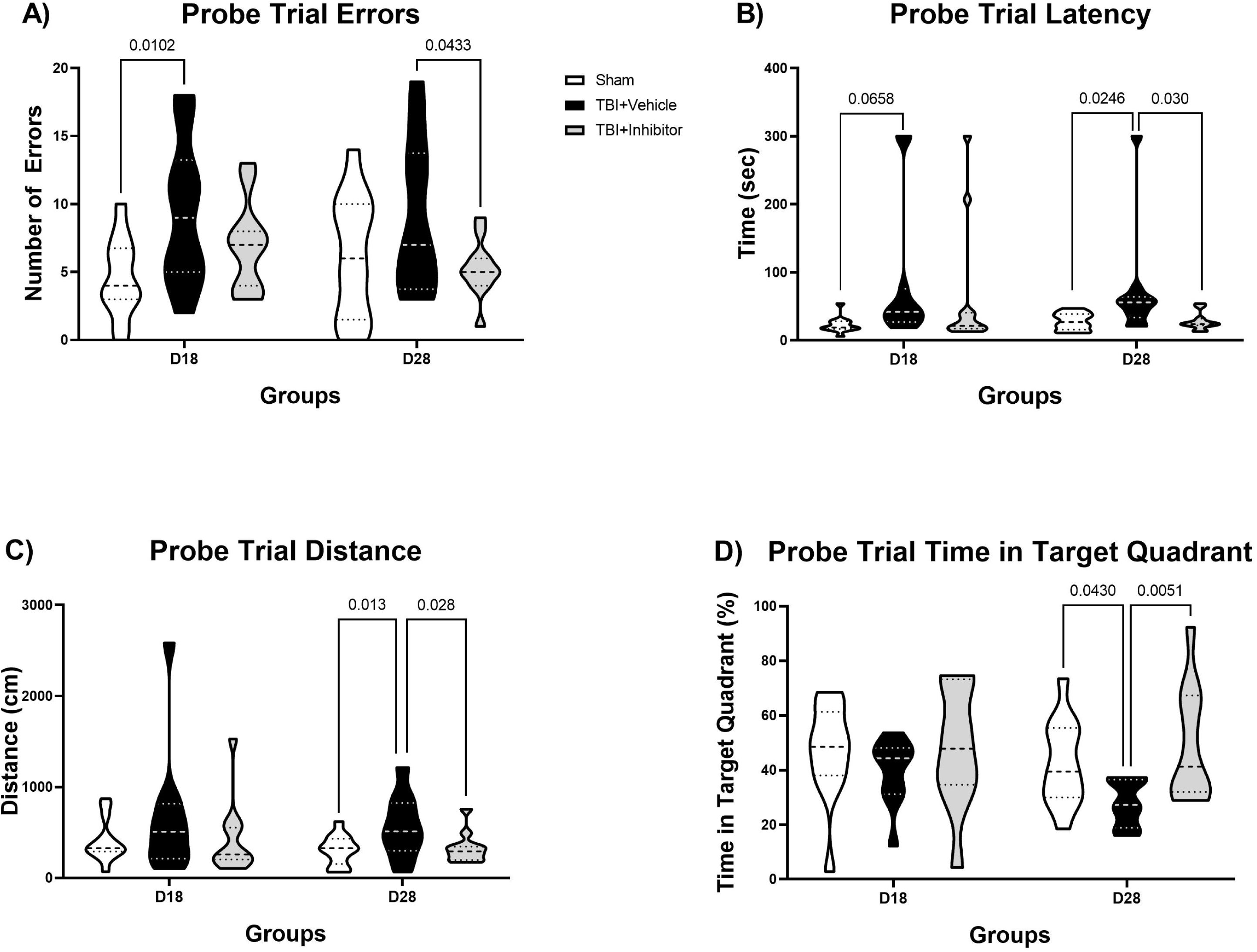
Violin plots of performance on the short-term (18 DPI) and long-term (28 DPI) memory probes of the Barnes maze after fluid percussion or sham injury. The TBI+Vehicle group made significantly more errors (A) and had a trend for longer latency to find the target hole (B) compared to the Sham group on 18DPI. On 28DPI, the TBI+Vehicle group had a significantly longer latency to the target hole (B), travelled a significantly longer distance to locate the target hole (C), and spent significantly less time in the target quadrant (D) compared to both the Sham and TBI+Inhibitor groups. There was also a trend for a higher number of errors on 28DPI in the TBI+Vehicle group compared to the TBI+Inhibitor group. Statistical comparisons were made with ordinary one-way ANOVAs and Fisher’s LSD for each parameter at each time point.

### Open Field Test

There were no significant differences between groups in the percent of time spent in the inner region of the open field or in the total distance travelled on D1 (Fig 5), although there was a trend for less time in the inner region in the TBI+Vehice group compared to the Sham group (*p*=0.08). On D25 there was a significant difference in the percent of time spent in the inner region (*F_2,32_*=5.71, *p*=0.0076), with the TBI+Vehicle group spending significantly less time in the inner region than both the sham and TBI+Inhibitor groups (*p*=0.04 vs Sham; *p*=0.002 vs TBI+Inhibitor). There were no differences in distance travelled on this day.

**Figure 5.**
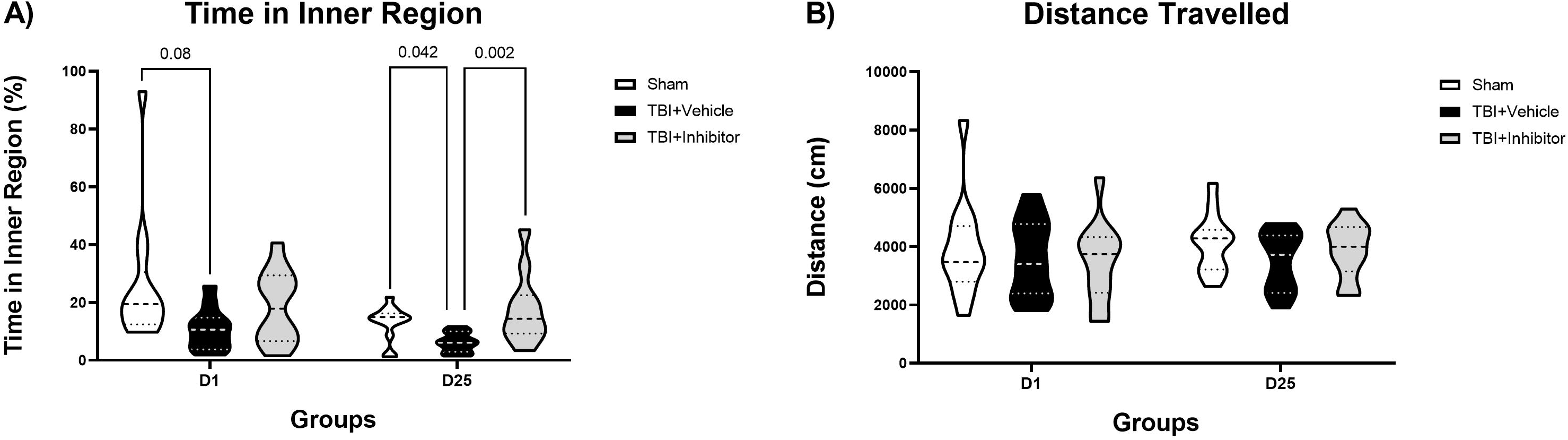
Violin plots of rat performance on the open field test one- and 25-days post-injury (DPI). There was a trend for less time spent in the inner region of the open field in the TBI+Vehicle compared with the Sham group at one DPI (A). The TBI+Vehicle group spend significantly less time than either the Sham or TBI+Inhibitor groups in the inner region of the open field on 25DPI (A). There were no differences in the distance travelled between any groups (B). Statistical comparisons were made with ordinary one-way ANOVAs and Fisher’s LSD for each parameter at each time point.

### Chronic KT ratios

The KT ratio was measured in the ipsilateral and contralateral perilesional neocortex and hippocampus at 28 days post injury, after behavioural testing was complete. There were no significant differences in the KT-ratio of any groups in either brain region (Fig. 1E, F).

## Discussion

The KP has been implicated in a number of neurological and psychiatric disorders (Zwilling et al., 2011; Vécsei et al., 2012; Lovelace et al., 2016; Savonije et al., 2023), including TBI in humans (Yan et al, 2015). Increases in neurotoxic metabolites of the KP have been associated with worse functional recovery (Chung et al., 2009; Zakhary et al., 2020; Meier et al., 2022). It is important to understand whether the KP is similarly upregulated in experimental models of TBI in order to determine whether manipulation of this pathway may be a therapeutic target to improve TBI outcomes. In this study we have shown increased IDO1 activity, as measured by an increased KT ratio, in the perilesional neocortex and hippocampus three days after FPI, with normalization by seven DPI. This increase in activity is blocked by oral administration of the IDO1 inhibitor PF-06840003, which also led to improved performance in measures of memory (Barnes maze) and anxiety-like behaviors (open field). These findings suggest IDO1 and the KP should be further investigated for their mechanistic roles in TBI outcomes and as potential treatment targets.

We have shown increased IDO1 activity as demonstrated by an increased ratio of kynurenine:tryptophan. IDO1 is the key enzyme driving the conversion of tryptophan to kynurenine in the kynurenine pathway during pathological conditions (Wirthgen et al., 2018). The increased KT ratio found three days after FPI is consistent with research showing increased IDO1 activity in highly inflammatory conditions (Badawy et al., 2017; Pallotta et al., 2021).

There is a significant neuroinflammatory response to FPI (Clausen et al., 2019; Ma et al., 2019; Peterson et al., 2015) and other experimental models of TBI (Ma et al., 2019) that can trigger changes in the KP by stimulating increased IDO1 activity (Yan et al., 2015; Zhang et al., 2018). This increased IDO1 activity can then lead to increased production of neurotoxic metabolites, such as QuinA, which have been implicated in worse outcomes after TBI in humans (Yan et al., 2015). Oral administration of the IDO1 inhibitor PF-06840003 was effective in preventing the increase in KT ratio three days after FPI. It is important to note that the first dose of PF-06840003 was only administered *after* FPI. This is very relevant clinically, as it demonstrates there is a window of time after FPI during which treatment can be initiated. Further studies will be needed to delineate the crucial time period for treatment. In addition, the effectiveness of an *oral* IDO1 inhibitor after FPI would be very beneficial in the clinical setting, avoiding the need for more invasive local or parenteral therapies. Oral PF-06840003 has already demonstrated a tolerable safety profile in humans (Reardon et al., 2020).

IDO1 inhibition with PF-06840003 prevented spatial memory deficits found after FPI, particularly at the later timepoint tested. While FPI did not impact rat learning ability on the Barnes maze at 14-17 DPI, it did cause memory disruption. The lack of learning deficits after FPI is in keeping with previous reports (Fitzgerald et al., 2022; Krishna et al., 2017; Lima, Camila Nayane de Carvalho et al., 2019) and not unexpected. FPI-induced memory deficits were not as marked on the short-term probe trial as they were on the long-term probe. TBI+Vehicle rats did however make significantly more errors than Sham rats before finding the target hole on 18 DPI, a pattern not seen in the TBI+Inhibitor group. There was a greater effect of FPI on the long-term memory probe trial at 28 DPI, with the TBI+Vehicle group having a significantly increased latency to the target hole, travelling a significantly greater distance before locating the target hole, and spending significantly less time in the target quadrant than either the Sham or TBI+Inhibitor groups. These results point to the involvement of IDO1 in the long-term visuospatial memory deficits seen after TBI. The hippocampus plays a key role in learning and recall of short-term and long-term visuospatial memory (Suarez et al., 2019), and increased IDO1 activity was found in the ipsilateral hippocampus after FPI in this study. The exact mechanism by which IDO1 inhibition improved memory outcomes after FPI is not known. It is possible that increased IDO1 activity after FPI resulted in a greater production of QuinA, which could damage and destroy hippocampal neurons. This reasoning would be consistent with research from Alzheimer’s studies which suggest that QuinA can directly cause excitotoxicity and neuronal cell damage in hippocampal cells, causing long-term spatial memory loss (Himanshu et al., 2020; Knight et al., 2021; Rahman et al., 2009). This may at least in part be mediated by increased oxidative stress (Mor et al., 2021). By excessively binding to NMDA receptors, QuinA causes constant influxes of calcium into neuronal cells which contribute to the production of reactive oxygen species and elevates the levels of oxidative stress in brain tissue (Mor et al., 2021). While our results show promise for IDO1 manipulation as a target to improve memory after TBI, the mechanisms remain to be elucidated.

FPI did not impact distance travelled in the open field early after injury, and there was only a trend for a decreased percent of time spent in the inner region at 1 DPI. However, the TBI+Vehicle group spent significantly less time in the inner region of the open field at 25 DPI compared to both the Sham and TBI+Inhibitor groups. This pattern of behavior is indicative of increased anxiety (Himanshu et al., 2020; Knight et al., 2021) and is consistent with reports of increased anxiety-like behaviors in rats after FPI demonstrated in a variety of tests (Bao et al., 2012; Fucich et al., 2019; Almeida-Suhett et al., 2014). Increased IDO1 activity has been linked with anxious behaviours in response to environmental stress (Lima, Camila Nayane de Carvalho et al., 2019), and IDO1 inhibition following an inflammatory challenge significantly reduced previously present anxious behaviours (Salazar et al., 2012). This may be related to the ability of QuinA to severely disrupt dopaminergic metabolism in the amygdala and mesolimbic system (Kurachi et al., 2000). Studies in athletes that have suffered sports related mild TBIs also found a significant increase in plasma QuinA levels which correlated with increased anxiety and depressive symptoms (Meier & Savitz, 2022; Singh et al., 2016). Our findings provide further support that IDO1 plays a role in mood disturbances after TBI and that IDO1 inhibition may help in prevention.

We have also shown a mild impact of IDO1 inhibition on motor recovery as assessed by the composite Neuroscore. While the TBI+Vehicle group continued to perform worse than its baseline at 28 DPI, a difference from baseline was no longer found at this timepoint for the TBI+Inhibitor group. However, there was no significant difference between these two groups at any time point. It is possible that treatment with PF-06840003 sped up the rate of recovery, but if assessments had been performed later than 28 DPI the overall improvement in the TBI+Vehicle group compared to its baseline may have matched that in the TBI+Inhibitor group. Longer-term studies would be helpful in this regard. However, no differences were found in the total distance travelled in the open field at either time point tested, another suggestion that gross motor function was not impacted. The composite Neuroscore assesses grip strength and motor reflexes in rats but does not accurately capture muscle strength, gait or specific limb impairments, therefore it would be important to assess outcome on other motor behavior tests before coming to a conclusion as to whether IDO1 inhibition impacts motor ability after TBI. However, it is possible that IDO1 may not play a significant role in the development of sensorimotor dysfunction following TBI.

The present study demonstrates increased IDO1 activity in a commonly used model of TBI, fluid percussion injury, and is proof of principle that IDO1 deserves further attention as a possible target to improve TBI outcomes. While we have shown improved performance on memory and anxiety-like behavioral tests, there are limitations to this study that will need to be addressed in future work. We have shown IDO1 activity is increased three DPI and normalizes by seven DPI, but it will be important to further characterize the temporal profile of TBI-induced changes in the KP in order to determine how large is the time window to begin intervention. Since downstream metabolites such as QuinA seem to be heavily implicated in the adverse effects of TBI, it is crucial that future studies also investigate the effects of IDO1 inhibition on KynA and QuinA levels. In addition, oral PF-06840003 was given for 28 DPI but it may not need to be dosed for so long. Only male rats were used in this study, but given the positive results described here ongoing studies are addressing the role of sex in KP-mediated outcomes after TBI. Sex differences play a role in both TBI (Nguyen et al., 2016; Valera et al., 2021; Gupte et al., 2019) and the KP (Meier et al., 2018; Fertan et al., 2019; Buck et al., 2020; Baratta et al., 2018), but there are no reports of sex differences in the KP in relation to TBI or in terms of IDO1 inhibition. Mechanistic studies will also be needed to determine how IDO1 influences TBI outcomes. The effects of IDO1 inhibition on neuroinflammation and oxidative stress after TBI are ripe areas for investigation.

This study supports a role for the KP and IDO1 in injury mechanisms and outcomes of experimental TBI. IDO1 activity is transiently increased after FPI and can be targeted to improve behavioral outcomes. As IDO1 activity is mainly a result of pathological conditions, this may be a helpful target to improve outcomes with minimal side effects after TBI. This proof of principle study demonstrates further work in this area is warranted.

## Author’s Contributions

Marawan Sadek contributed to the conceptualization, formal analysis, investigation, methodology, writing, review, and editing.

Kurt R Stover contributed to the conceptualization, investigation, methodology, review, and editing.

Xiaojing Liu contributed to the methodology, review, and editing. Donald F Weaver contributed to the resources, review, and editing.

Mark A Reed contributed to the conceptualization, resources, review, and editing.

Aylin Y Reid contributed to the conceptualization, formal analysis, funding acquisition, methodology, project administration, supervision, validation, writing, review, and editing.

## Acknowledgements

The authors would like to thank Chiping Wu and Mingdong Yang for their technical assistance with the work in this study. We would also like to thank the Canadian Institutes for Health Research, the Ontario Graduate Scholarship fund, and the University of Toronto for their financial support of this project.

## Literature Cited

Ahlstedt, J., Konradsson, E., Ceberg, C., & Redebrandt, H. N. (2020). Increased effect of two-fraction radiotherapy in conjunction with IDO1 inhibition in experimental glioblastoma. PloS One, 15(5), e0233617.

Almeida-Suhett, C. P., Prager, E. M., Pidoplichko, V., Figueiredo, T. H., Marini, A. M., Li, Z., . ..Braga, M. F. M. (2014). Reduced GABAergic inhibition in the basolateral amygdala and the development of anxiety-like behaviors after mild traumatic brain injury. PLoS ONE, 9(7), e102627.

Badawy, A. A.-B. (2017). Kynurenine pathway of tryptophan metabolism: Regulatory and functional aspects. International Journal of Tryptophan Research, 10, 117864691769193.

Bao, F., Shultz, S. R., Hepburn, J. D., Omana, V., Weaver, L. C., Cain, D. P., & Brown, A. (2012). A CD11d monoclonal antibody treatment reduces tissue injury and improves neurological outcome after fluid percussion brain injury in rats. Journal of Neurotrauma, 29(14), 2375–2392.

Baratta AM, Buck SA, Buchla AD, Fabian CB, Chen S, Mong JA, Pocivavsek A (2018). Sex Differences in Hippocampal Memory and Kynurenic Acid Formation Following Acute Sleep Deprivation in Rats. Sci Rep. 8(1):6963.

Buck SA, Baratta AM, Pocivavsek A (2020). Exposure to elevated embryonic kynurenine in rats: Sex-dependent learning and memory impairments in adult offspring. Neurobiol Learn Mem. 174:107282.

Casillas-Espinosa PM, Andrade P, Santana-Gomez C, Paananen T, Smith G, Ali I, Ciszek R, Ndode-Ekane XE, Brady RD, Tohka J, Hudson MR, Perucca P, Braine EL, Immonen R, Puhakka N, Shultz SR, Jones NC, Staba RJ, Pitkänen A, O’Brien TJ. Harmonization of the pipeline for seizure detection to phenotype post-traumatic epilepsy in a preclinical multicenter study on post-traumatic epileptogenesis. Epilepsy Res. 2019 Oct;156:106131. doi: 10.1016/j.eplepsyres.2019.04.011. Epub 2019 Apr 27. PMID: 31076256; PMCID: PMC6814157.

Cheong, J. E., Ekkati, A., & Sun, L. (2018). A patent review of IDO1 inhibitors for cancer. Expert Opinion on Therapeutic Patents, 28(4), 317–330.

Chung, R. S., Leung, Y. K., Butler, C. W., Chen, Y., Eaton, E. D., Pankhurst, M. W., . . . Guillemin, G. J. (2009). Metallothionein treatment attenuates microglial activation and expression of neurotoxic quinolinic acid following traumatic brain injury. Neurotoxicity Research, 15(4), 381–389.

Clausen, F., Marklund, N., & Hillered, L. (2019). Acute inflammatory biomarker responses to diffuse traumatic brain injury in the rat monitored by a novel microdialysis technique. Journal of Neurotrauma, 36(2), 21–211.

Crosignani S, Bingham P, Bottemanne P, Cannelle H, Cauwenberghs S, Cordonnier M, Dalvie D, Deroose F, Feng JL, Gomes B, Greasley S, Kaiser SE, Kraus M, Négrerie M, Maegley K, Miller N, Murray BW, Schneider M, Soloweij J, Stewart AE, Tumang J, Torti VR, Van Den Eynde B, Wythes M. Discovery of a Novel and Selective Indoleamine 2,3-Dioxygenase (IDO-1) Inhibitor 3-(5-Fluoro-1H-indol-3-yl)pyrrolidine-2,5-dione (EOS200271/PF-06840003) and Its Characterization as a Potential Clinical Candidate. J Med Chem. 2017 Dec 14;60(23):9617–9629. doi: 10.1021/acs.jmedchem.7b00974. Epub 2017 Nov 21. PMID: 29111717.

Dobos, N., de Vries, Erik F. J, Kema, I. P., Patas, K., Prins, M., Nijholt, I. M., . . . Eisel, U. L. M. (2012). The role of indoleamine 2,3-dioxygenase in a mouse model of neuroinflammation-induced depression. Journal of Alzheimer’s Disease, 28(4), 905–915.

Dolšak, A., Gobec, S., & Sova, M. (2021). Indoleamine and tryptophan 2,3-dioxygenases as important future therapeutic targets. Pharmacology & Therapeutics, 221, 107746.

Febinger, H., Thomasy, H., & Gemma, C. (2016). A controlled cortical impact mouse model for mild traumatic brain injury. Bio-Protocol, 6(16).

Fertan E, Stover KRJ, Brant MG, Stafford PM, Kelly B, Diez-Cecilia E, Wong AA, Weaver DF, Brown RE (2019). Effects of the Novel IDO Inhibitor DWG-1036 on the Behavior of Male and Female 3xTg-AD Mice. Front Pharmacol. 10:1044.

Fitzgerald, J., Houle, S., Cotter, C., Zimomra, Z., Martens, K. M., Vonder Haar, C., & Kokiko-Cochran, O. N. (2022). Lateral fluid percussion injury causes sex-specific deficits in anterograde but not retrograde memory. Frontiers in Behavioral Neuroscience, 16, 806598.

Fucich, E. A., Mayeux, J. P., McGinn, M. A., Gilpin, N. W., Edwards, S., & Molina, P. E. (2019). A novel role for the endocannabinoid system in ameliorating motivation for alcohol drinking and negative behavioral affect after traumatic brain injury in rats. Journal of Neurotrauma, 36(11), 1847–1855.

Gomes, B., Driessens, G., Bartlett, D., Cai, D., Cauwenberghs, S., Crosignani, S., Dalvie, D., Denies, S., Dillon, C. P., Fantin, V. R., Guo, J., Letellier, M.-C., Li, W., Maegley, K., Marillier, R., Miller, N., Pirson, R., Rabolli, V., Ray, C., Kraus, M. (2018). Characterization of the selective indoleamine 2,3-dioxygenase-1 (IDO1) catalytic inhibitor EOS200271/PF-06840003 supports IDO1 as a critical resistance mechanism to pd-(l)1 blockade therapy. Molecular Cancer Therapeutics, 17(12), 2530–2542.

Gupte R, Brooks W, Vukas R, Pierce J, Harris J. (2019) Sex Differences in Traumatic Brain Injury: What We Know and What We Should Know. J Neurotrauma. Jul 19. doi: 10.1089/neu.2018.6171.

Harrison, F. E., Reiserer, R. S., Tomarken, A. J., & McDonald, M. P. (2006). Spatial and nonspatial escape strategies in the barnes maze. Learning & Memory (Cold Spring Harbor, N.Y.), 13(6), 809–819.

Hattori, S., Takao, K., Funakoshi, H., & Miyakawa, T. (2018). Comprehensive behavioral analysis of tryptophan 2,3-dioxygenase (*tdo2*) knockout mice. Neuropsychopharmacology Reports, 38(2), 52–60.

Himanshu, Dharmila, Sarkar, D., & Nutan. (2020). A review of behavioral tests to evaluate different types of anxiety and anti-anxiety effects. Clinical Psychopharmacology and Neuroscience : The Official Scientific Journal of the Korean College of Neuropsychopharmacology, 18(3), 341–351.

Kabadi, S. V., Hilton, G. D., Stoica, B. A., Zapple, D. N., & Faden, A. I. (2010). Fluid-percussion–induced traumatic brain injury model in rats. Nature Protocols, 5(9), 1552–1563.

Knight, P., Chellian, R., Wilson, R., Behnood-Rod, A., Panunzio, S., & Bruijnzeel, A. W. (2021). Sex differences in the elevated plus-maze test and large open field test in adult wistar rats. Pharmacology, Biochemistry and Behavior, 204, 173168.

Kovesdi, E., Kamnaksh, A., Wingo, D., Ahmed, F., Grunberg, N. E., Long, J. B., Agoston, D. V. (2012). Acute minocycline treatment mitigates the symptoms of mild blast-induced traumatic brain injury. Frontiers in Neurology, 3, 111.

Krishna, G., Agrawal, R., Zhuang, Y., Ying, Z., Paydar, A., Harris, N. G., . . . GomezPinilla, F. (2017). 7,8-dihydroxyflavone facilitates the action exercise to restore plasticity and functionality: Implications for early brain trauma recovery. Biochimica Et Biophysica Acta, 1863(6), 1204–1213.

Kurachi, M., Sumiyoshi, T., Shibata, R., Sun, Y., Uehara, T., Tanii, Y., & Suzuki, M. (2000). Changes in limbic dopamine metabolism following quinolinic acid lesions of the left entorhinal cortex in rats. Psychiatry and Clinical Neurosciences, 54(1), 83–89.

Learoyd, A. E., & Lifshitz, J. (2012). Comparison of rat sensory behavioral tasks to detect somatosensory morbidity after diffuse brain-injury. Behavioural Brain Research, 226(1), 197–204.

Lima, Camila Nayane de Carvalho, da Silva, Francisco Eliclécio Rodrigues, Chaves Filho, Adriano José Maia, Queiroz, Ana Isabelle de Gois, Okamura, Adriana Mary Nunes Costa, Fries, G. R., Macedo, D. S. (2019). High exploratory phenotype rats exposed to environmental stressors present memory deficits accompanied by immune-inflammatory/oxidative alterations: Relevance to the relationship between temperament and mood disorders. Frontiers in Psychiatry, 10, 547.

Liu, Y., Li, S., Gao, Z., Li, S., Tan, Q., Li, Y., Wang, D., & Wang, Q. (2021). Indoleamine 2,3-dioxygenase 1 (IDO1) promotes cardiac hypertrophy via a PI3K-akt-mtordependent mechanism. Cardiovascular Toxicology, 21(8), 655–668.

Lovelace MD, Varney B, Sundaram G, Franco NF, Ng ML, Pai S, Lim CK, Guillemin GJ, Brew BJ. Current Evidence for a Role of the Kynurenine Pathway of Tryptophan Metabolism in Multiple Sclerosis. Front Immunol. 2016 Aug 4;7:246. doi: 10.3389/fimmu.2016.00246. PMID: 27540379; PMCID: PMC4972824.

Lynch, K., Gradecki, S., Kwak, M., Meneveau, M., Wages, N., Gru, A., & Slingluff, C. (2020). IDO1 expression in melanoma metastases is low and associated with improved overall survival. The American Journal of Surgical Pathology, 45(6), 787–795.

Ma, X., Aravind, A., Pfister, B. J., Chandra, N., & Haorah, J. (2019). Animal models of traumatic brain injury and assessment of injury severity. Molecular Neurobiology, 56(8), 5332–5345.

Maegele, M., Lippert-Gruener, M., Ester-Bode, T., Sauerland, S., Schäfer, U., Molcanyi, M., . . . Neugebauer, E. A. M. (2005). Reversal of neuromotor and cognitive dysfunction in an enriched environment combined with multimodal early onset stimulation after traumatic brain injury in rats. Journal of Neurotrauma, 22(7), 772–782.

Mazarei, G., & Leavitt, B. R. (2015). Indoleamine 2,3 dioxygenase as a potential therapeutic target in huntington’s disease. Journal of Huntington’s Disease, 4(2), 109–118.

McIntosh, T. K., Vink, R., Noble, L., Yamakami, I., Fernyak, S., Soares, H., & Faden, A. L. (1989). Traumatic brain injury in the rat: Characterization of a lateral fluid-percussion model. Neuroscience, 28(1), 233–244.

Meier TB, Drevets WC, Teague TK, Wurfel BE, Mueller SC, Bodurka J, Dantzer R, Savitz J (2018). Kynurenic acid is reduced in females and oral contraceptive users: Implications for depression. Brain Behav Immun. 67:59–64.

Meier, T. B., & Savitz, J. (2022). The kynurenine pathway in traumatic brain injury: Implications for psychiatric outcomes. Biological Psychiatry (1969), 91(5), 449–458.

Mor, A., Tankiewicz-Kwedlo, A., Krupa, A., & Pawlak, D. (2021). Role of kynurenine pathway in oxidative stress during neurodegenerative disorders. Cells (Basel, Switzerland), 10(7), 1603.

Ndode-Ekane, X. E., Santana-Gomez, C., Casillas-Espinosa, P. M., Ali, I., Brady, R. D., Smith, G., Andrade, P., Immonen, R., Puhakka, N., Hudson, M. R., Braine, E. L., Shultz, S. R., Staba, R. J., O’Brien, T. J., & Pitkänen, A. (2019). Harmonization of lateral fluid-percussion injury model production and post-injury monitoring in a preclinical multicenter Biomarker Discovery Study on post-traumatic epileptogenesis. Epilepsy Research, 151, 7–16.

Nguyen R, Fiest KM, McChesney J, Kwon CS, Jette N, Frolkis AD, Atta C, Mah S, Dhaliwal H, Reid A, Pringsheim T, Dykeman J, Gallagher C. (2016) The international incidence of traumatic brain injury: a systematic review and meta-analysis. Can J Neurol Sci. 43:774–785.

O’Brien, T. J., Cardamone, L., Liu, Y. R., Hogan, R. E., Maccotta, L., Wright, D. K., Zheng, P., Koe, A., Gregoire, M.-C., Williams, J. P., Hicks, R. J., Jones, N. C., Myers, D. E., Shultz, S. R., & Bouilleret, V. (2013). Can structural or functional changes following traumatic brain injury in the rat predict epileptic outcome? Epilepsia, 54(7), 1240–1250.

Pallotta, M. T., Rossini, S., Suvieri, C., Coletti, A., Orabona, C., Macchiarulo, A., Volpi, C., & Grohmann, U. (2021). Indoleamine 2,3-dioxygenase 1 (Ido1): An up-to-date overview of an eclectic immunoregulatory enzyme. The FEBS Journal, 289(20), 6099–6118.

Peterson, T. C., Maass, W. R., Anderson, J. R., Anderson, G. D., & Hoane, M. R. (2015). A behavioral and histological comparison of fluid percussion injury and controlled cortical impact injury to the rat sensorimotor cortex. Behavioural Brain Research, 294, 254–263.

Pitkänen, A., Immonen, R. J., Gröhn, O. H. J., & Kharatishvili, I. (2009). From traumatic brain injury to posttraumatic epilepsy: What animal models tell us about the process and treatment options. Epilepsia, 50(s2), 21–29.

Rahman, A., Ting, K., Cullen, K. M., Braidy, N., Brew, B. J., & Guillemin, G. J. (2009). The excitotoxin quinolinic acid induces tau phosphorylation in human neurons. PLoS ONE, 4(7), e6344.

Reardon, D. A., Desjardins, A., Rixe, O., Cloughesy, T., Alekar, S., Williams, J. H., Li, R., Taylor, C. T., & Lassman, A. B. (2020). A phase 1 study of Pf-06840003, an oral indoleamine 2,3-dioxygenase 1 (IDO1) inhibitor in patients with recurrent malignant glioma. Investigational New Drugs, 38(6), 1784–1795.

Reid, A. Y., Bragin, A., Giza, C. C., Staba, R. J., & Engel, J. (2016). The progression of electrophysiologic abnormalities during epileptogenesis after experimental traumatic brain injury. Epilepsia, 57(10), 1558–1567.

Rowe, R. K., Harrison, J. L., O’Hara, B. F., & Lifshitz, J. (2014). Recovery of neurological function despite immediate sleep disruption following diffuse brain injury in the mouse: Clinical relevance to medically untreated concussion. Sleep (New York, N.Y.), 37(4), 743–752.

Salazar, A., Gonzalez-Rivera, B. L., Redus, L., Parrott, J. M., & O’Connor, J. C. (2012). Indoleamine 2,3-dioxygenase mediates anhedonia and anxiety-like behaviors caused by peripheral lipopolysaccharide immune challenge. Hormones and Behavior, 62(3), 202–209.

Saletti PG, Mowrey WB, Liu W, Li Q, McCullough J, Aniceto R, Lin IH, Eklund M, Casillas-Espinosa PM, Ali I, Santana-Gomez C, Coles L, Shultz SR, Jones N, Staba R, O’Brien TJ, Moshé SL, Agoston DV, Galanopoulou AS; EpiBioS4Rx Study Group. Early preclinical plasma protein biomarkers of brain trauma are influenced by early seizures and levetiracetam. Epilepsia Open. 2023 Jun;8(2):586–608. doi: 10.1002/epi4.12738. Epub 2023 Apr 25. PMID: 37026764; PMCID: PMC10235584.

Savitz, J. (2020). The kynurenine pathway: A finger in every pie. Molecular Psychiatry, 25(1), 131–147.

Savonije K, Meek A, Weaver DF. Indoleamine 2,3-Dioxygenase as a Therapeutic Target for Alzheimer’s Disease and Geriatric Depression. Brain Sci. 2023 May 24;13(6):852. doi: 10.3390/brainsci13060852. PMID: 37371332; PMCID: PMC10296628

Singh, R., Savitz, J., Teague, T. K., Polanski, D. W., Mayer, A. R., Bellgowan, P. S. F., & Meier, T. B. (2016). Mood symptoms correlate with kynurenine pathway metabolites following sports-related concussion. Journal of Neurology, Neurosurgery and Psychiatry, 87(6), 670–675.

Sinz, E. H., Kochanek, P. M., Heyes, M. P., Wisniewski, S. R., Bell, M. J., Clark, R. S. B., . . . Marion, D. W. (1998). Quinolinic acid is increased in CSF and associated with mortality after traumatic brain injury in humans. Journal of Cerebral Blood Flow and Metabolism, 18(6), 610–615.

Suarez, A. N., Noble, E. E., & Kanoski, S. E. (2019). Regulation of memory function by feeding-relevant biological systems: Following the breadcrumbs to the hippocampus. Frontiers in Molecular Neuroscience, 12, 101.

Tian, C., Chen, L., Chen, H., Huan, X., Hu, J., Shen, J., Miao, Z. (2019). Inhibition of the BET family reduces its new target gene IDO1 expression and the production of L-kynurenine. Cell Death & Disease, 10(8), 557–13.

Valera EM, Joseph ALC, Snedaker K, Breiding MJ, Robertson CL, Colantonio A, Levin H, Pugh MJ, Yurgelun-Todd D, Mannix R, Bazarian JJ, Turtzo LC, Turkstra LS, Begg L, Cummings DM, Bellgowan PSF. (2021). Understanding traumatic brain injury in females: a state-of-the-art summary and future directions. J Head Trauma Rehabil. 36(1):E1–E17.

Vécsei, L., Szalárdy, L., Fülöp, F., & Toldi, J. (2012). Kynurenines in the CNS: Recent advances and new questions. Nature Reviews Drug Discovery, 12(1), 64–82.

Wirthgen, E., Hoeflich, A., Rebl, A., & Günther, J. (2018). Kynurenic acid: The janus-faced role of an immunomodulatory tryptophan metabolite and its link to pathological conditions. Frontiers in Immunology, 8.

Wu, W., Nicolazzo, J. A., Wen, L., Chung, R., Stankovic, R., Bao, S. S., . . . Guillemin, G. J. (2013). Expression of tryptophan 2,3-dioxygenase and production of kynurenine pathway metabolites in triple transgenic mice and human alzheimer’s disease brain. PLoS ONE, 8(4), e59749.

Yan, E. B., Frugier, T., Lim, C. K., Heng, B., Sundaram, G., Tan, M., Morganti Kossmann, M. C. (2015). Activation of the kynurenine pathway and increased production of the excitotoxin quinolinic acid following traumatic brain injury in humans. Journal of Neuroinflammation, 12(1), 110.

Zakhary, G., Sherchan, P., Li, Q., Tang, J., & Zhang, J. H. (2020). Modification of kynurenine pathway via inhibition of kynurenine hydroxylase attenuates surgical brain injury complications in a male rat model. Journal of Neuroscience Research, 98(1), 155–167.

Zhang, C., Saatman, K. E., Royo, N. C., Soltesz, K. M., Millard, M., Schouten, J. W., . . . McIntosh, T. K. (2005). Delayed transplantation of human neurons following brain injury in rats: A long-term graft survival and behavior study. Journal of Neurotrauma, 22(12), 1456–1474

Zhang, Z., Rasmussen, L., Saraswati, M., Koehler, R. C., Robertson, C., & Kannan, S. (2018). Traumatic injury leads to inflammation and altered tryptophan metabolism in the juvenile rabbit brain. Journal of Neurotrauma, 36(1), 74–86.

Zwilling, D., Huang, S.-Y., Sathyasaikumar, K. V., Notarangelo, F. M., Guidetti, P., Wu, H.-Q., Lee, J., Truong, J., Andrews-Zwilling, Y., Hsieh, E. W., Louie, J. Y., Wu, T., Scearce-Levie, K., Patrick, C., Adame, A., Giorgini, F., Moussaoui, S., Laue, G., Rassoulpour, A., & Muchowski, P. J. (2011). Kynurenine 3-monooxygenase inhibition in blood ameliorates neurodegeneration. Cell, 145(6), 863–874.

